# Air monitoring by nanopore sequencing

**DOI:** 10.1101/2023.12.19.572325

**Authors:** Tim Reska, Sofya Pozdniakova, Sílvia Borràs, Michael Schloter, Lídia Cañas, Albert Perlas Puente, Xavier Rodó, Yuanyuan Wang, Barbro Winkler, Jörg-Peter Schnitzler, Lara Urban

**Affiliations:** Helmholtz AI, Helmholtz Zentrum München, Neuherberg, Germany; Helmholtz Pioneer Campus, Helmholtz Zentrum München, Neuherberg, Germany; Technical University of Munich, School of Life Sciences, Freising, Germany; AIRLAB, Climate and Health (CLIMA) group, ISGlobal, Barcelona, Spain; Institute of Comparative Microbiome Analysis, Helmholtz Zentrum München, Neuherberg, Germany; Catalan Institution for Research and Advanced Studies, ICREA, Barcelona, Spain; Technical University of Munich, School of Engineering and Design, Munich, Germany; Research Unit Environmental Simulation (EUS), Helmholtz Zentrum München, Neuherberg, Germany

**Author notes:** contributed equally (Tim Reska, Sofya Pozdniakova).

## Abstract

While the air microbiome and its diversity are essential for human health and ecosystem resilience, comprehensive air microbial diversity monitoring has remained rare, so that little is known about the air microbiome’s composition, distribution, or functionality. Here we show that nanopore sequencing-based metagenomics can robustly assess the air microbiome in combination with active air sampling through liquid impingement and tailored computational analysis. We provide fast and portable laboratory and computational approaches for air microbiome profiling, which we leverage to robustly assess the taxonomic composition of the core air microbiome of a controlled greenhouse environment and of a natural outdoor environment. We show that long-read sequencing can resolve species-level annotations and specific ecosystem functions through *de novo* metagenomic assemblies despite the low amount of fragmented DNA used as an input for nanopore sequencing. We then apply our pipeline to assess the diversity and variability of an urban air microbiome, using Barcelona, Spain, as an example; this randomized experiment gives first insights into the presence of highly stable location-specific air microbiomes within the city’s boundaries, and showcases the robust microbial assessments that can be achieved through automatable, fast, and portable nanopore sequencing technology.

## Introduction

The air microbiome encompasses a broad spectrum of bioaerosols, including bacteria, archaea, fungi, viruses, bacterial endotoxins, mycotoxins, and pollen (1). While its pivotal functions for human health and ecosystem resilience are recognized, little is known about its composition, distribution, and functionality (2). Past research efforts, particularly those driven by infectious diseases such as COVID-19 and tuberculosis, have shifted the research focus towards potentially pathogenic microbial taxa; however, exposure to a diverse air microbiome has also been increasingly considered as a health-promoting factor, underscoring the need for holistic air microbial diversity monitoring (3).

Most genetics-based air microbiome studies have employed targeted sequencing via metabarcoding due to the low biomass of bioaerosols (1, 4). While metabarcoding increases the sensitivity of taxonomic detection, it is inherently limited by amplification biases and incomplete databases. In contrast, metagenomics, which is based on shotgun sequencing of native DNA, avoids amplification biases and allows for *de novo* reconstructions of microbial genomes for robust species identification and functional annotation. Such metagenomic approaches have also recently been applied for low biomass bioaerosol analysis (7) and have revealed the complex nature and diverse origins of the air microbiome (4), including vertical-altitudinal stratification of microbial abundance and distribution (5), and substantial diurnal, seasonal, temperature-, and humidity-dependent fluctuations (6).

These metagenomic assessments of the air microbiome have thus far relied on short-read sequencing technology, which provides accurate sequencing data but hampers *de novo* assemblies, especially of highly repetitive genomic regions, and accurate species- or strain-level identification due to the inherently short sequencing reads; long-read sequencing, on the other hand, has facilitated *de novo* genome assemblies (8) and assessments of highly repetitive genomic regions, including the detection of antimicrobial resistance genes (9), from metagenomic data. Especially recent advances in nanopore sequencing technology have made long-read sequencing increasingly relevant for microbial diversity assessments due to the technology’s substantially improving sequencing accuracy ([ref](https://pubmed.ncbi.nlm.nih.gov/35789207/); [ref](https://www.biorxiv.org/content/10.1101/2024.03.15.585313v2)) while maintaining its long-read sequencing capacity and its automatable (10), fast, and portable deployability for applications in clinical (11) or remote settings (12). While nanopore sequencing has been used to characterize the microbial diversity of various environments, such as of freshwater (13) and dust (14), no approaches have yet been established to leverage the technology’s unique advantages for monitoring the taxonomic and functional diversity of the air microbiome.

Here, we established laboratory and computational approaches to enable robust air microbiome profiling through nanopore metagenomics. We first evaluated the suitability of long-read shotgun sequencing for assessing the air microbiome in a controlled indoor environment, and then applied our approaches to an outdoor environment for validation. We showed that nanopore sequencing is a robust tool to describe the composition and diversity of microbial taxa in the air, and to concurrently annotate *de novo* microbial genomes to evaluate potential human health consequences. We finally applied our laboratory and computational approaches to conduct a randomized air sampling campaign in Barcelona, Spain, to robustly describe its urban air microbiome.

## Materials and Methods

We first conducted preliminary tests to compare standard air sampling and DNA extraction approaches for nanopore sequencing-based air metagenomics; this included the testing of standard quartz filter- and liquid impingement-based air samplers and the optimization of respective DNA extraction approaches for subsequent nanopore shotgun sequencing, which relies on minimum DNA input without nucleotide amplification and is sensitive to native DNA contamination (**Supplementary Information**: *Air sampling and DNA extraction optimizations*).

Based on these preliminary tests, we decided to use the Coriolis μ liquid impinger (Bertin Instruments, France; (**Supplementary Information**: *Air sampling and DNA extraction optimizations*) for air sampling, which uses cyclonic forces to concentrate airborne biomass into a collection liquid in a cone. We used 15 mL of ultrapure water with 0.005 % Triton-X (Sigma-Aldrich, Germany) as collection liquid, which functions as a nonionic surfactant to enhance organic compound solubility and surface enlargement due to foam generation. The liquid impinger was positioned at 1.5 m above the ground to sample air within the human breathable zone, which ranges from 1.4 to 1.8 m. We operated the liquid impinger at an air flow rate of 300 L min^-1^ and at a collection liquid refilling rate of 0.8 mL min^-1^ to counter liquid evaporation during sampling. After sampling, we directly transferred the collected liquid into a sterile 15 mL falcon tube. We then divided the liquid across three 5 mL tubes, centrifuged them at 18,000 x g for 25 min, and collected the pellets. The pellets were resuspended, aggregated, and subsequently centrifuged twice at 18,000 x g for 25 min while discarding the supernatant.

We first sampled air in a greenhouse (“Gh”; Helmholtz Munich Environmental Research Unit) as a controlled environment with moderate human activity and continuous air circulation (mean ambient temperature of 23 °C); we sampled air for three consecutive days, either for 1h in three consecutive replicates per day or for 3h with one replicate per day (**Supplementary Table 1**). We next sampled air in a natural environment (“Nat”), namely on the Helmholtz Munich campus on the outskirts of Munich (48.220889, 11.597028), which is mainly surrounded by natural grassland. We sampled for six consecutive days, following an alternating pattern of 3h or 6h of air sampling; we here tested 6h as sampling duration since we expected a higher variability in the air microbiome in comparison to the controlled greenhouse setting (**Supplementary Table 1**). The liquid impinger was positioned in a shaded area to avoid significant thermal fluctuations. While the weather remained relatively constant and sunny across the six sampling days (ambient temperature ranged from 21°C to 25°C, and humidity from 42% to 71%.), we note that the 6h-sample from day 4 was affected by rain and thunderstorm at the end of the sampling activity. We finally collected urban air samples in Barcelona, Spain, from October 16th to November 3rd, 2023. We sampled five different urban locations: Gracia (“Residential Area”, 41.398861, 2.153490), Eixample (“City Center”, 41.385500, 2.155103), Poblenou (“Urban Beach”, 41.404135, 2.206550), Vall d’Hebron (“Outer Belt”, 41.425887, 2.148349), and Observatori Fabra (“Green Belt”, 41.419772, 2.122447). We conducted randomized sampling in terms of timing (morning *versus* afternoon) and across days; each location was sampled three times for 3h using two Coriolis μ air samplers, respectively, resulting in altogether 30 air samples (**Supplementary Table 1**).

**Table 1.**
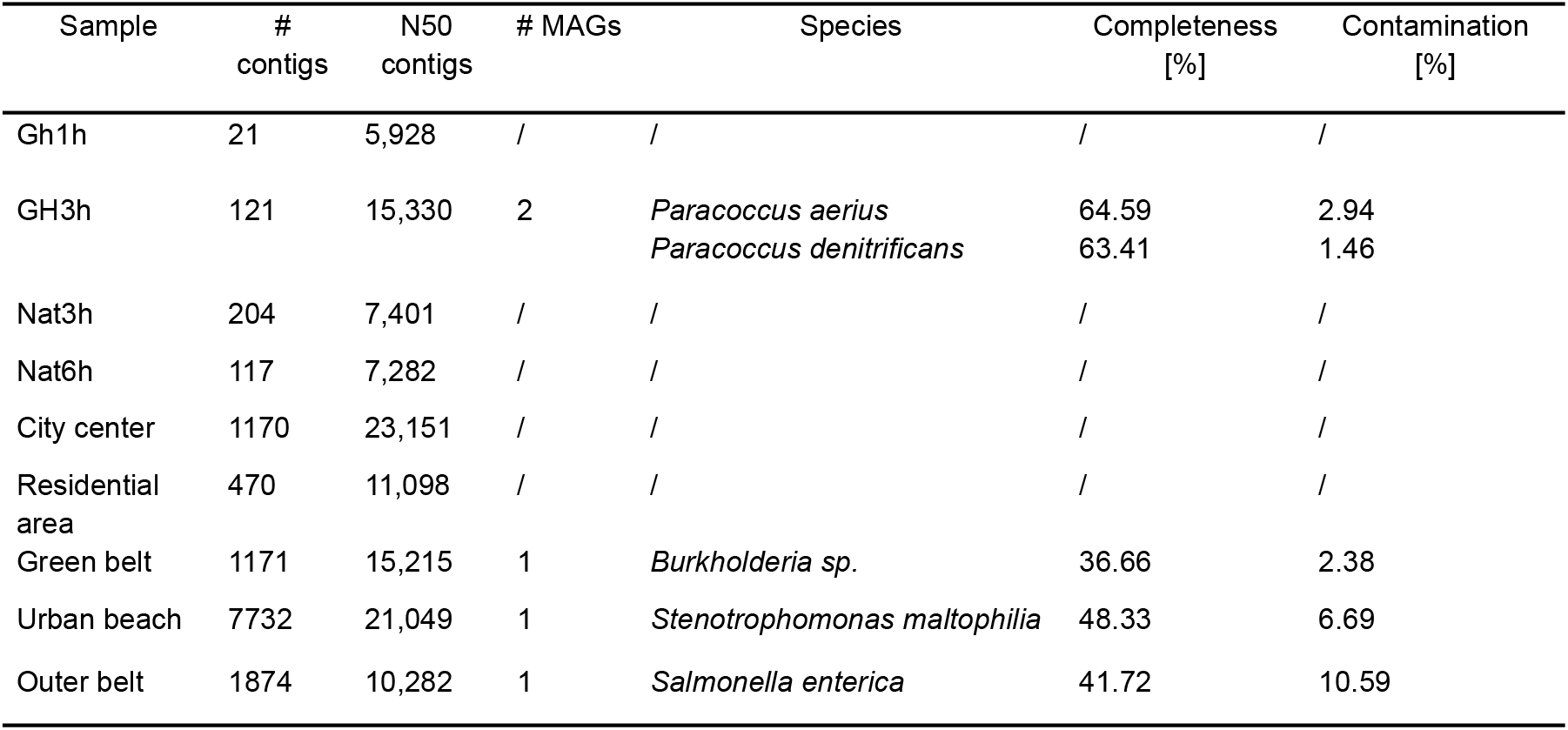
*De novo* genome assembly results across all air samples from the controlled (Gh), natural (Nat) environment, and the urban microbiome dataset. Contigs were assembled and then used to identify metagenome-assemblies (MAGs), their taxonomic origin, completeness, and contamination.

Based on our preliminary tests, we further decided to use the spin-column based PowerSoil Pro Kit (QIAGEN, 2018, Hilden, Germany) for DNA extractions, using 30 μL of elution buffer (**Supplementary Information**: *Air sampling and DNA extraction optimizations*). Final DNA concentration was measured on a Qubit 4.0 fluorometer (Invitrogen, 2021), using the high-sensitivity DNA kit and 3 μL of DNA elution as input per sample. We then used the Rapid Barcoding library preparation kit (RBK114-24 V14), R10.4.1 MinION flow cells, and MinKNOW by Oxford Nanopore Technologies (Oxford, UK) to nanopore shotgun-sequence the extracted DNA of the air samples. During library preparation, we used each barcode twice per air sample to increase the DNA input per sample. For sequencing the samples of the controlled and natural environment, we used one R10.4.1 flow cell per sample type (i.e., for all 1h-, 3h-, or 6-samples and replicates, respectively). For sequencing the samples of the urban environment, we pooled all samples from the Outer Belt location onto one flow cell (since they exhibited the lowest DNA concentrations), and the samples of the City Center and Residential Area, as well as of the Green Belt and Urban Beach, onto one flow cell, respectively. The sequencing parameters included a minimum read length of 20 bases, a translocation speed of 400 bases per second, and each sequencing run lasted 24 hours. As we used MinKNOW v23.04.3 for the controlled and natural environment, this sequencing data was generated at a signal measurement frequency of 4 kHz, whereas we used the updated MinKNOW v23.04.5 for the urban environment, which generated sequencing data at 5 kHz.

We included negative controls along our entire protocol to identify contamination of the low-biomass air samples. For sampling negative controls, we treated one liquid impinger cone per sampling event the same way that we treated the actual sampling cone, but we only left them in the impinger for a few minutes and did not actively sample air. For the urban environment, negative sampling controls were collected once per sampling day and sampling location. For DNA extraction and sequencing negative controls, we included one sample of 700 μL nuclease-free water (Thermo Fisher Scientific) per DNA extraction and one sample of 20 μL nuclease-free water per library preparation, respectively. We barcoded all negative controls, i.e. sampling, extraction, and sequencing controls, and included them in the same sequencing library as the respective control samples. We further subjected a positive control of five Gram-positive bacteria, three Gram-negative bacteria, and two fungal species (ZymoBIOMICS Microbial Community Standard, D6300) to our DNA extraction and sequencing protocols to assess any potential biases. The positive control was sequenced on a separate flow cell since the high DNA concentration would have outcompeted the low-biomass air samples.

We next used Guppy v6.3.2 (r10.4.1_e8.2_400bps_hac; [ref](https://community.nanoporetech.com/docs/prepare/library_prep_protocols/Guppy-protocol/v/gpb_2003_v1_revax_14dec2018/guppy-software-overview)) in high-accuracy (HAC) mode for basecalling the controlled and natural environment samples, and Dorado v4.3.0 (dna_r10.4.1_e8.2_400bps_hac@v4.3.0; [ref](https://github.com/nanoporetech/dorado)) for HAC-basecalling of the urban environment samples. We only processed the data that had passed internal data quality thresholds during sequencing (“passed” sequencing reads). Porechop v0.2.3 ([ref](https://github.com/rrwick/Porechop)) was used for removing sequencing adapters and barcodes, and Nanofilt v2.8.0 ([ref](https://academic.oup.com/bioinformatics/article/34/15/2666/4934939)) was applied for filtering reads at a minimum average quality score of 8 and a minimum length of 100 bases for all samples. We then used Kraken2 v2.0.7 ([ref](https://genomebiology.biomedcentral.com/articles/10.1186/s13059-019-1891-0)) with the NCBI nt database (access 29.01.2023) for taxonomic classification across all samples, and downsampled them to a specific read count for comparable taxonomic assessments across samples of one sample type: 5k reads for 1h-samples from the controlled environment, 15k reads for the 3h-samples from the controlled environment, 70k reads for the natural environment samples, and 30k reads for the urban environment samples. We performed Principal Coordinate Analysis (PCoA) on the relative abundances of the genera identified in the urban environment samples, which were downsampled to 30k read, using Python v3.9 with Pandas v1.3.3, NumPy v1.21.2, scikit-learn v0.24.2, scikit-bio v0.5.6, SciPy v1.7.1, and Matplotlib v3.5.2.. The 20 most abundant microbial genera at a minimum relative abundance of 1% as well as the PCoA were visualized using matplotlib v3.5.2 in Python v3.9. We additionally benchmarked several additional bioinformatic analysis tool in application to the controlled and natural environment samples, including DIAMOND BLASTX ([ref](https://www.nature.com/articles/nmeth.3176)) for protein-based taxonomic classifications and the Chan-Zuckerberg (CZID) computational pipeline ([ref](https://academic.oup.com/gigascience/article/9/10/giaa111/5918865)) for hybrid taxonomic classifications (i.e., as a combination of read- and contig-based classification).

We generated *de novo* assemblies using metaflye v2.9.1 ([ref](https://www.nature.com/articles/s41592-020-00971-x)), followed by polishing with minimap2 v2.17 ([ref](https://academic.oup.com/bioinformatics/article/34/18/3094/4994778)) and three rounds of Racon v1.5 ([ref](https://www.ncbi.nlm.nih.gov/pmc/articles/PMC5411768/)). The resulting contigs were then binned into Metagenome-Assembled Genomes (MAGs) using metaWRAP v1.3 ([ref](https://microbiomejournal.biomedcentral.com/articles/10.1186/s40168-018-0541-1)), which integrates the output of various binning tools. The MAGs were refined and quality-checked using CheckM v1.2.2 ([ref](https://github.com/Ecogenomics/CheckM/wiki/Overview)). We only maintained MAGs at minimum completeness of 30% and maximum contamination of 10%. For the urban microbiome dataset, we pooled across all samples per sampling location to maximize the number of reads before binning. We finally applied functional annotation to our metagenomic dataset to assess the presence of general metabolic pathways and ecosystem functions (**Supplementary Information**: *Functional annotation*); to identify antimicrobial resistance and virulence genes, we applied AMRFinderPlus v3.12.8 ([ref](https://www.nature.com/articles/s41598-021-91456-0)) and ABRicate v1.0.1 ([ref](https://github.com/tseemann/abricate)) to the reads, contigs, and bins; for the application to the read level, we converted the fastq files to fasta files using seqkit v2.8.2 ([ref](https://www.ncbi.nlm.nih.gov/pmc/articles/PMC5051824/)).

To obtain information about the anthropogenic impact on the different urban sampling locations, we obtained remote sensing data (Sentinel-2 L1C orthoimage products from October 24th 2023) that provides top-of-atmosphere reflectance, which we used to classify the city of Barcelona into Local Climate Zones (LCZs) on based ten bands with 10 m and 20 m ground sampling distances ([ref](https://www.sciencedirect.com/science/article/pii/S0034425719304912)). We further used the portable aerosol spectrometer Dust Decoder 11-D (GRIMM Aerosol Technik GmbH, Germany) to monitor particle mass fractions (TSP, PM_10_, and PM_2.5_; TSP=total suspended particles; PM=particulate matter) as well as temperature and relative humidity measurements in 1-minute intervals during each sampling event. We then summarized and analyzed the resulting data using Python v3.9 and SciPy v1.13.0: We applied the Kruskal-Wallis and post-hoc Dunn’s tests to identify significant environmental differences between locations, and conducted regression analyses to assess correlations between particle mass fractions and microbial diversity indices (Shannon, Simpson, and richness of microbial genera).

## Results

After confirming that Coriolis μ liquid impingement resulted in sufficient high-quality DNA yield for nanopore shotgun sequencing after one hour of sampling (**Materials and Methods**; **Supplementary Information**: *Air sampling and DNA extraction optimizations*), we conducted a pilot study in a controlled environment to determine the robustness of the metagenomic data and assess the impact of sampling duration (**Materials and Methods**). For the 1h-samples, DNA yields ranged from 17.7 ng to 50.7 ng (0.98 ng/m^3^ to 2.82 ng/m^3^), while the 3-hour samples showed DNA yields ranging from 130.2 ng to 179.4 ng (2.41 ng/m^3^ to 3.32 ng/m^3^; **Supplementary Table 1**; *pilot_study* sheet). Nanopore shotgun sequencing delivered between 7k and 60k high-quality sequencing read at a median read length of 896 bases (**Figure 1A**), respectively, of which 5k to 35k reads were successfully mapped to the taxonomic genus level using Kraken2 and the NCBI nt database (**Figure 1B-C**; **Supplementary Table 1**; *pilot_study* sheet). After downsampling to the same number of reads per sample type (1h- and 3h-samples, respectively), the taxonomic composition of the 20 most abundant taxa indicated that only the 3-hour sampling duration captured a stable “core” air microbiome across days at the genus level (**Figure 1D-E**). These assessments were consistent for protein-level or hybrid read- and assembly-based methods, both at the taxonomic phylum and genus level (**Supplementary Figures 1-2)**. The most abundant genera included soil- and plant-associated bacteria such as *Bradyrhizobium, Paracoccus, Nocardioides, Massilia*, and *Streptomyces* (**Figure 1D-E**; **Materials and Methods**).

**Figure 1.**
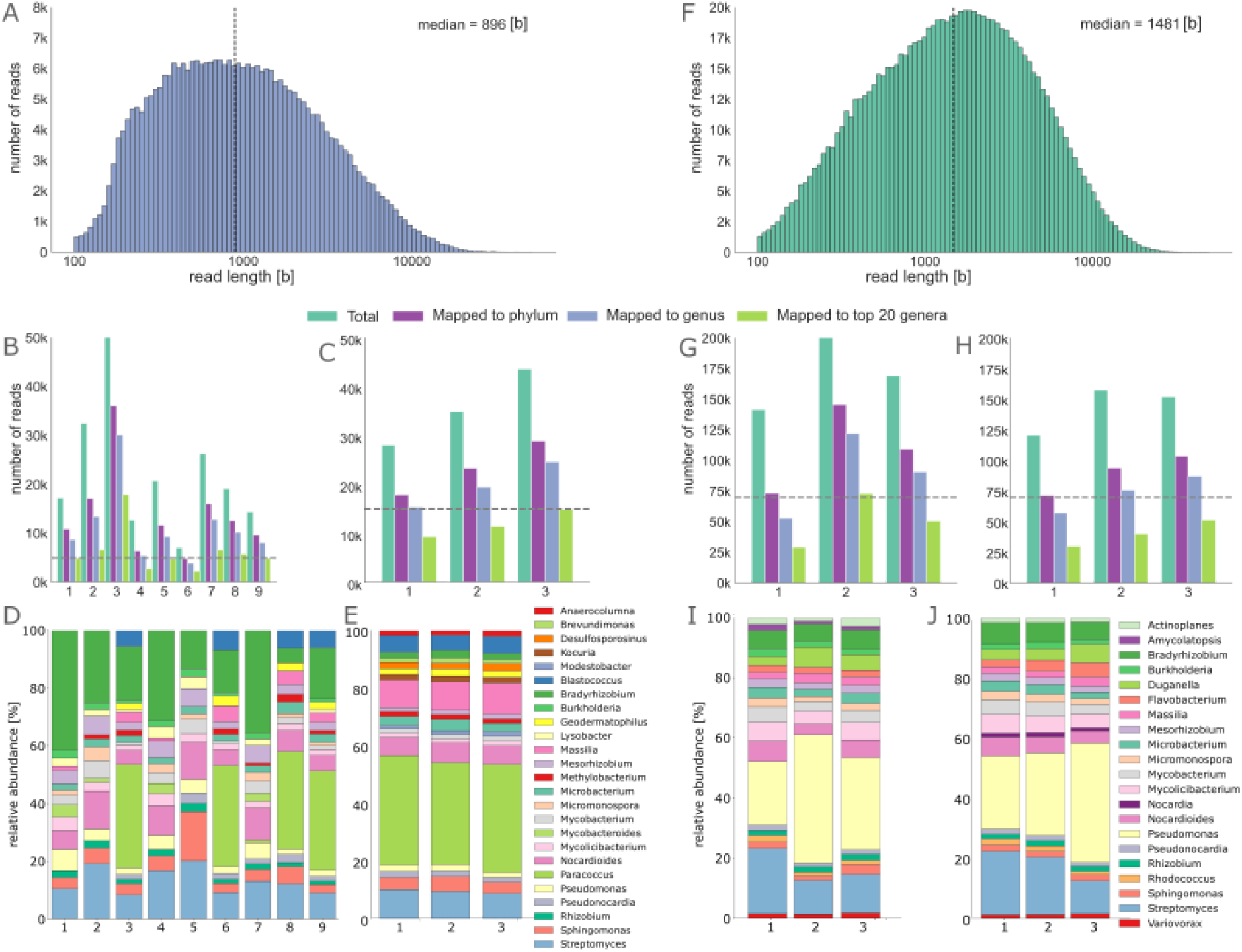
Robust air microbiome assessments of a controlled (*left*; **A-E**) and natural (*right*; **F-J**) environment through nanopore shotgun sequencing. **A**. Nanopore sequencing read length distribution across 1h- and 3-Gh samples. Number of total sequencing reads, and of reads mapping to taxonomic phylum and genus level as well as to the 20 most dominant genera using Kraken2 (Material and methods) across the **B**. 1h- and **C**. 3-Gh samples; the downsampling threshold across samples is indicated by the dashed horizontal line. Taxonomic composition of the **D**. 1h- and **E**. 3h-samples after downsampling based on the 20 most dominant genera across samples. **F**. Air microbiome monitoring in a natural environment (Nat). **G**. Nanopore sequencing read length distribution across 3h- and 6h-Nat samples. Number of total sequencing reads, and of reads mapping to taxonomic phylum and genus level as well as to the 20 most dominant genera using Kraken2 across the **H**. 3h- and **J**. 6-Gh samples; the downsampling threshold across samples is indicated by the dashed horizontal line. Taxonomic composition of the **I**. 3h- and **J**. 6h-samples after downsampling based on the 20 most dominant genera across samples.

Based on these results, we conducted a pilot study in a natural environment over six days; we sampled air for either 3h or 6h, assuming that the natural environment might show more variability than the controlled environment and require longer sampling duration. Briefly, while the extended sampling duration increased total DNA yield, it did not consistently increase the amount of biomass per cubic meter of sampled air, suggesting diminishing returns in efficiency with longer durations (**Supplementary Table 1**; *pilot_study* sheet). Nanopore shotgun sequencing resulted 130k to 200k high-quality sequencing reads at a slightly higher median read length than the controlled environment of 1481 (**Figure 1F**), of which 70k to 140k reads were successfully mapped to the taxonomic genus level (**Figure 1G-H**; **Supplementary Table 1**; *pilot_study* sheet). After downsampling all samples to 70k reads, analysis of the relative abundance of 20 most abundant taxa revealed a very similar profile for both 3-hour and 6-hour samples. The taxonomic assignments were again consistent across protein-level or hybrid read- and assembly-based methods, both at the taxonomic phylum and genus level (**Supplementary Figures 1-2)**. A distinct air microbiome profile was observed in the natural environment in comparison to the controlled settings, with high predominance of *Pseudomonas* and unique detection of microbial taxa such as *Actinoplanes, Amycolatopsis, Dugnaella, Flavobacterium, Nocardia, Rhodococcus*, and *Variovorax* (**Figure 1I-J**; **Materials and Methods**).

All negative controls resulted in low DNA yields (of <0.1 ng) from typical contaminant species such as *Escherichia, Salmonella, Shigella, Francisella*, and *Pseudomonas* (**Supplementary Figure 3A-B**; **Material and Methods**) (17). This demonstrates that no external contamination had influenced our assessment of air as a low-biomass ecosystem, thus underscoring the reliability of the presented results. The application of our protocol to a well-defined mock community further showed that all bacterial and fungal species could be detected with approximately correct abundance estimates. Although the fungal taxa and Gram-positive *Bacillus subtilis*, in particular, were underrepresented (**Supplementary Figure 3C**; **Material and Methods**).

We finally applied our optimized laboratory and computational approaches to assess an exemplary urban microbiome using nanopore metagenomics (**Figure 2A**; *left*; **Materials and Methods**). Our remote-sensing-based Local Climate Zones (LCZs) (**Figure 2A**; *right*) indicated that most of our sampling locations (City Center, Residential Area, and Urban Beach) were of the compact low-rise category, a typical feature of central urban environments. The Outer Belt location was classified as compact mid-rise category, which features taller buildings at the outskirts of the city. The Green Belt location was classified as scattered trees category, featuring more natural elements. In terms of air pollution assessed through particle mass fractions (**Supplementary Table 2**; **Materials and Methods**), we found significant differences in TSP, PM_10_ and PM_2.5_, between our sampling locations (**Supplementary Figure 4**). The total air pollution measured by TSP was highest in the three compact low-rise sampling locations, while TSP was lowest in the Outer Belt. The relatively medium levels of TSP in the Green Belt were dominated by relatively high levels of PM_10_ (**Supplementary Figure 4**).

**Figure 2.**
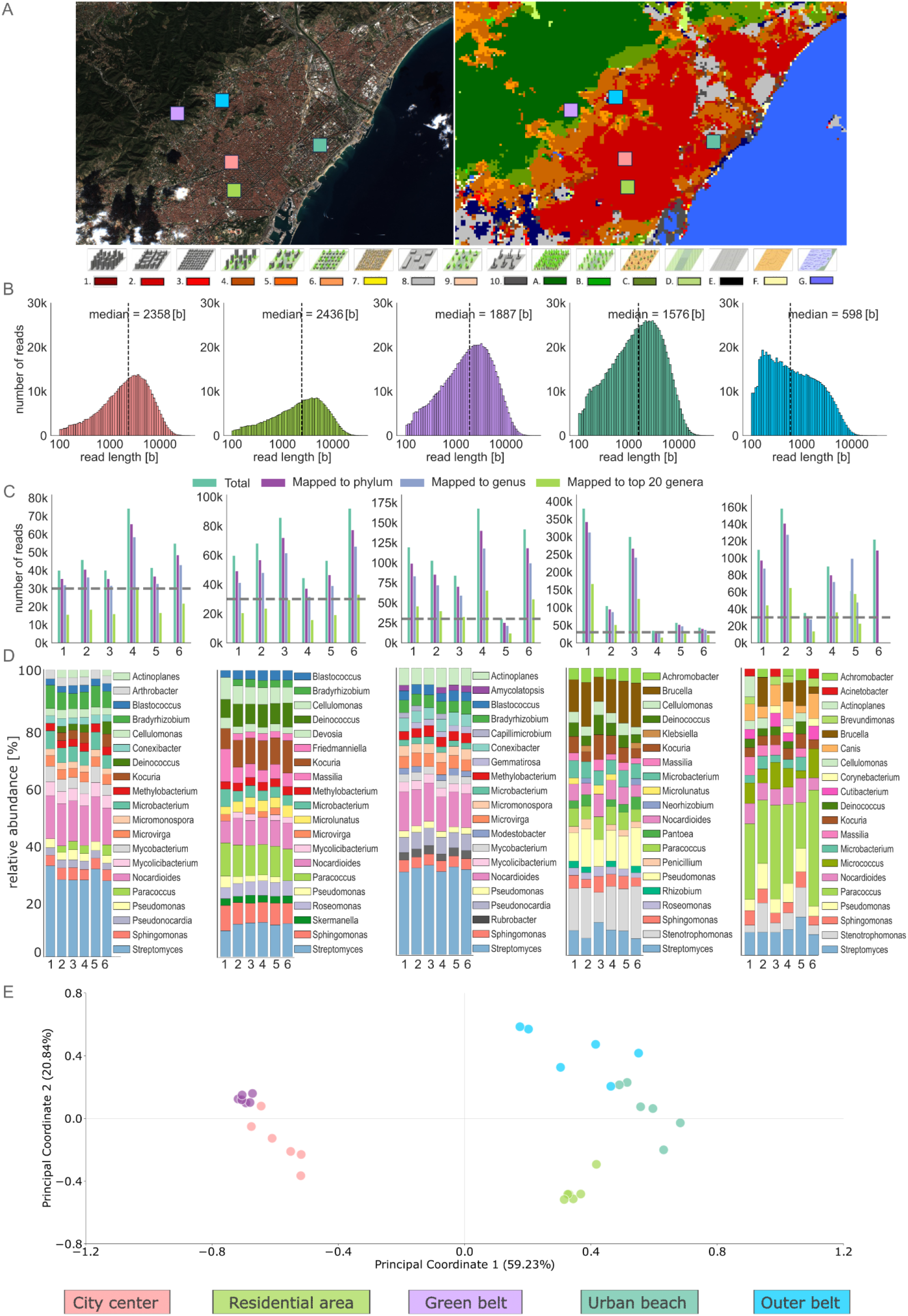
Local climate zones classification and metagenomic analysis of Barcelona using satellite images and nanopore shotgun sequencing data. **A**. Sentinel-2 image and LCZs classification map of Barcelona on 24.10.2023, with a legend of LCZs classes at the bottom. The colored squares indicate the five sampling locations. The legend at the bottom depicts various Local Climate Zones (LCZs) represented by 3D models and their corresponding colors. (1-Compact high-rise; 2-Compact mid-rise; 3-Compact low-rise; 4-Open high-rise; 5-Open mid-rise; 6-Open low-rise; 7-Lightweight low-rise; 8-Large low-rise; 9-Sparsely built; 10-Heavy industry; A-Dense trees; B-Scattered trees; C-Bush scrub; D-Low plants; E-Bare rock or paved; F-Bare soil or sand; G-Water). **B**. Histograms showing the distribution of read lengths [b] for each sampling site, with the median read length indicated on the top of each histogram. **C**. Bar plots displaying the number of reads mapped at various taxonomic levels for each sample site, with the downsampling threshold indicated by the dashed horizontal line. **D**. Relative abundance of the top 20 most abundant bacterial genera at the read level, downsampled to 30K reads before taxonomic classification using Kraken2. **E**. Principal Coordinates Analysis (PCoA) of the relative abundances of the bacterial genera identified at the five sampling locations.

Nanopore shotgun sequencing delivered between 33k and 422k high-quality sequencing read at a median read length of between 598 and 2358 bases (**Figure 2B**), respectively, of which 21k to 312k reads were successfully mapped to the taxonomic genus level using Kraken2 and the NCBI nt database (**Figure 2C**; **Supplementary Table 1**; *urban_study* sheet). The City Center exhibited the longest DNA fragments, and the Outer Belt location the shortest DNA fragments (**Figure 2B**). The relatively high fragmentation in the Outer Belt coincided with generally low DNA yields across all the location’s samples and replicates (**Supplementary Table 1**; *urban_study* sheet).

For taxonomic comparisons across replicates and samples, we again downsampled the number of reads (here to 30k reads per sample) and compared the relative distribution of the 20 most abundant microbial genera per location at a minimum relative abundance cutoff of 1% displaying (**Materials and Methods**). We observed that the microbial compositions were highly location-specific across all six samples per location, including across the three randomized sampling events and the two respective sampling replicates (**Figure 2D**; **Materials and Methods**). The core urban air microbiome consisted of microbial genera such as *Streptomyces, Sphingamonas, Pseudomonas, Nocardioides*, and *Microbacterium*, which were detected across all samples. Specifically the Green Belt was characterized by the presence of several unique taxa such as *Rubrobacter, Gemmatirosa, Capillimicrobium*, and *Amycolatopsis*, whereas dominant “urban” taxa such as *Paracoccus, Kocuria, Deinociccus*, and *Cellulomonas* were not detected at all (**Figure 2D**). Principal Coordinate Analysis (PCoA) clearly distinguishes the five different urban locations, with the first PCoA axis separating the Green Belt and City Center locations from the remaining ones; the second PCoA axis then further delineates the individual sampling locations (**Figure 2E**).

Despite the location-specific differences in air microbial composition (**Figure 2A**), in LCZ-based land usage (**Figure 2D**) and in air pollution measured by particle mass fractions (**Supplementary Figure 4**), we found no significant correlations between any environmental variable and microbial diversity measurements (**Materials and Methods**).

To next obtain as highly contiguous *de novo* genome assemblies as possible, we pooled all samples per location before contig assembly and binning (**Materials and Methods**). Taxonomic classification of these bins showed that only the most abundant taxa could be assembled (**Table 1**). Functional annotation of the reads, contigs, and bins detected typical microbial metabolic functions (**Supplementary Information**: *Functional annotation*). We next focused on the annotation of antimicrobial resistance and virulence genes with potential human health consequences (**Supplementary Table 3**; **Materials and Methods**). One of the most frequently detected genes was the *VanR-O* gene, which is responsible for vancomycin resistance. When comparing resistance gene prevalence across urban locations, the Urban Beach location exhibited the highest density of resistance genes; the *blaCARB-8* and *blaCARB-16* genes, which confer beta-lactam resistance, and the *blaOXA-17* gene, which confers oxacillin resistance, were detected at the read level. Additionally, *blaL1*, which confers to a broad range of beta-lactam antibiotics, the *blaOXY* gene, which confers oxacillin resistance, and the *blaPSZ* gene, which confers resistance to penicillins and cephalosporins, were identified at the contig level (**Supplementary Table 3**).

## Discussion

Metagenomic approaches have provided unprecedented insights into the nature, origin, and complexity of the air microbiome (see Intro ref: currently 4-7). While past studies have relied on traditional short-read sequencing, we here describe the first long-read nanopore sequencing technology-based approaches to robustly assess the air microbiome. Although nanopore sequencing has been applied to various environmental samples, such as water and soil ([ref](https://www.ncbi.nlm.nih.gov/pmc/articles/PMC5755585/),[ref](https://elifesciences.org/articles/61504),[ref](https://elifesciences.org/articles/84553)), its applicability to air samples was expected to pose a particular challenge due to the ultra-low biomass of air and the amplification-free nature of nanopore sequencing ([ref](http://doi.org/10.1038/s41522-021-00209-4)). We here showed that nanopore shotgun sequencing in combination with active air sampling through liquid impingement and tailored computational analyses can reproducibly describe the air microbiome of different environments (**Figure 1**) while leveraging the latest nanopore chemistry improvements which offer high sequencing accuracy and reduced minimum DNA input requirements (see Intro refs: [ref;](https://pubmed.ncbi.nlm.nih.gov/35789207/) [ref](https://www.biorxiv.org/content/10.1101/2024.03.15.585313v2)).

We further showed that only three hours of active air sampling resulted in robust air microbiome assessments in a controlled and natural environment, with consecutive application of our laboratory and computational approaches to the urban air microbiome in Barcelona, Spain, revealing surprisingly stable location-specific signatures of microbial composition and diversity (**Figure 2**). These stable signatures could importantly be identified across replicates (using two air samplers per sampling event) and despite stringent randomization across sampling days and morning and afternoon sampling events. Several microbial taxa such as *Sphingomonas* and *Streptomyces*, which are known for their evolutionary adaptability, were nevertheless present in all air microbiomes, and could potentially be part of the stable air microbiome of this urban environment. Ordination of the taxonomic composition was able to capture the majority of variance in this multidimensional data (>80%; **Figure 2E**) and nicely visualizes the distinct clusters that separate each urban location and specifically the Green Belt and City Center locations from the remaining ones. The relative similarity of Green Belt and City Center samples might be attributable to the phenomenon of orographic uplift, where air masses ascend from lower regions (here the Barcelona City Center) to higher elevated areas (here the closeby Green Belt). As a result of this upward movement, certain airborne particles and microorganisms might have been transported from the City Center to the Green Belt location.

The individual samples of the Green Belt location cluster together most tightly (**Figure 2E**). This might be because of several microbial taxa that were uniquely detected at this location, which represents the only natural environment in our study according to our remote-sensing-based assessments; those unique taxa are known to be associated with soil or have been frequently found in forests and green spaces ([ref](https://www.ncbi.nlm.nih.gov/pmc/articles/PMC5485800/)). Besides this finding, we however found no evidence of correlation of the urban air microbiome with measurements of anthropogenic impact (as assessed through the remote-sensing-based Local Climate Zones, LCSz; **Figure 2A**) or of air pollution (as assessed through particle mass fraction measurements; **Supplementary Figure 4**). This might be due to complex interactions between air microbiomes, as exemplified by our hypothesis of the impact of orographic uplift, or because of lack of depth when describing our environmental variables. For example, air pollution by TSP was higher in the Green Belt than in the Outer Belt, which would have not been expected according to the remote-sensing-based anthropogenic impact inferences. However, these elevated levels of TSP in the Green Belt might have originated from natural air components such as pollen, which would require more in-depth environmental monitoring to dissect.

The annotation of antimicrobial resistance and virulence genes in our metagenomic data shows that we can use the same dataset to assess potential anthropogenic impacts on microbial diversity while concurrently understanding potential public health consequences ([ref](https://www.embopress.org/doi/full/10.15252/msb.202311686)). We detected evidence of antimicrobial resistance across all sampled environments (**Supplementary Table 3**), but especially the detections of clinically relevant beta-lactamases such as *blaCARB-8, blaOXA-1*, and *blal-1*, and of genes conferring resistance to other antibiotics such as carbenicillin and oxacillin ([ref](https://www.sciencedirect.com/science/article/pii/S0269749121009842)), in Barcelona’s urban air microbiome underscore the possibility of monitoring airborne virulence dissemination using nanopore-based metagenomics. Given the presence of *Brucella*, a typical canine pathogen, in several of our air samples, we further investigated our taxonomic annotation, which was based on the entire NCBI nt database, and were indeed able to detect the presence of *Canis lupus familiaris* in the same air samples. While this might point to a potential impact of animal domestication and specifically frequent dog walking in Barcelona on public health ([ref](https://www.nc.cdc.gov/eid/article/24/8/17-1171_article)), such interdependencies would have to be investigated in a controlled and/or experimental setting.

While we were able to build *de novo* assemblies from our nanopore-based air metagenomic data, most of the Metagenome-Assembled Genomes (MAGs) were incomplete (<30%) and/or showed high levels of contamination (>10%) (**Table 1**). Given the low amount of DNA input and therefore relatively small size of the resulting metagenomic datasets in combination with the expectedly high fragmentation of DNA in air samples, this might just be an inherent shortcoming when it comes to assessing the air microbiome – albeit applying long-read sequencing technology. We here found a particular small median DNA fragment and sequencing read length for the Outer Belt location (**Figure 2B**), which might point towards the impact of environmental conditions or specific taxonomic compositions (and variables such as the microorganisms’ genome size and cell wall composition) on the final fragment and read length distribution. It is further expected that non-viable microorganisms, which might significantly contribute to the air microbiome, result in more fragmented DNA in the air samples; this means that substantial differences in read lengths between microbial taxa might also be attributed to their differential viability in the air environment – a hypothesis that we might be able to resolve in the future using viability-resolved metagenomic approaches (cite Urel et al., 2024),

We emphasize that our sampling, laboratory and computational approaches constitute one feasible and reproducible way of using nanopore shotgun sequencing to profile the air microbiome. While we tested some additional established air sampling and DNA extraction methodology, we have not conducted an extensive study of all possible approaches. We specifically emphasize that the detection of fungi and Gram-positive bacteria could be improved when using different sample processing and DNA extraction techniques. This is also reflected by the application of our approaches to a positive control, which shows that fungal taxa and Gram-positive *Bacillus subtilis*, in particular, were underrepresented. As sturdier cell walls would require more aggressive DNA extraction approaches, this would, however, also lead to increased DNA fragmentation, especially in Gram-negative bacteria, and therefore more difficult downstream analyses. A good trade-off could be the sequencing of several, differently processed DNA extracts and subsequent data pooling to assess the microbial diversity of any air sample more holistically.

In conclusion, our study establishes a robust framework for air microbiome assessments using nanopore metagenomics. We envision that nanopore sequencing for air monitoring can provide a basis for fast, robust, and automated characterizations of the air microbiome in both urbanized and remote settings. This characterization importantly extends beyond taxonomic composition to include functions related to human and ecosystem health, such as pathogen and drug resistance and virulence gene detection, which can enhance our understanding of infectious disease transmission patterns and their relationship with exerted anthropogenic pressures.

## Data and code availability

All raw data has been made publicly available via ENA (study accession number: PRJEB76446). All code has been made publicly available via Github: https://github.com/ttmgr/Air_Metagenomics.

## Author contributions

T.R, S.P., S.B. and L.U. designed the experiment. T.R, S.P. conducted the sampling, processing, and data analysis under L.U.’s supervision. T.R, S.P., and L.U. wrote the manuscript with the contribution of all authors who revised and approved the manuscript.

## Supplementary Information

**Supplementary Table 1.** Sampling, DNA, and sequencing data of all air samples (*pilot_study* and *urban_study* sheets): sampling data includes date, temperature, humidity, and sampling duration; DNA metrics include total yield and yield per m^3^; and sequencing data includes total number of reads, filtered reads, read length distribution (median and N50), read-based taxonomic classification results using Kraken2, and assembly statistics using metaflye (Materials and Methods).

**Supplementary Table 2.** Environmental pollution of urban sampling location measured through particle mass fractions (TSP, PM10, and PM2.5; TSP=total suspended particles; PM=particulate matter); measurements were taken in one-minute intervals (Materials and Methods).

**Supplementary Table 3.** Antimicrobial resistance and virulence genes detected by ABRicate and AMRFinderPlus across all air samples (*pilot_study* and *urban_study* sheets) with respective gene coverage and mapping accuracy metrics (Materials and Methods).

### Air sampling and DNA extraction optimizations

We first tested two standard air sampling approaches, the high-volume sampler (HVS, MCV, Spain) and the Coriolis μ liquid impinger (Bertin Technologies, France), to assess the optimal air sampler for their compatibility with nanopore shotgun sequencing.

For the HVS, we used quartz filters for air sampling for 24h at a rate of 500 L/min. We applied both, phenol-chloroform extraction (16) and the standard PowerSoil Pro kit (QIAGEN, 2018), to the filters. While the phenol-chloroform method resulted in a higher total DNA yield than the standard extraction kit (data not shown), the nanodrop nucleic acid 260/280 measurements of around 1.2 indicated that the extracted DNA was highly contaminated, most likely due to residual phenol, which would block the nanopores during shotgun sequencing. The DNA yield of the standard extraction kit, on the other hand, was not sufficient for nanopore shotgun sequencing, which made us hypothesize that the standard kit – since not optimized for DNA extractions from quartz filter – might have chemically enhanced binding of the particles to the silica-enriched filters, and might therefore have made extraction inefficient.

For the liquid impingement-based and therefore filter-free sampler Coriolis μ, we sampled air for 1h at a rate of 300 L/min, and extracted sufficient DNA using the standard Qiagen kit: To increase DNA concentrations, we benchmarked that the volume of the final elution buffer (EB) could be reduced from the standard of 50 μL to 30μL. We further tested if a repeated washing of the spin column would further increase the DNA yield, which was not the case:

| Sample # | Volume EB | DNA concentration (ng/μL) | Total DNA (ng) |
| --- | --- | --- | --- |
| 1 | 50 μL | 0.131 | 6.55 |
| 2 | 50 μL | 0.283 | 14.15 |
| 3 | 50 μL | 0.330 | 16.50 |
| 4 | 30 μL | 1.240 | 37.2 |
| 5 | 30 μL | 0.767 | 23.01 |
| 6 | 30 μL | 1.69 | 50.7 |
| 7 | 30 μL twice | 0.607 | 18.09 |
| 8 | 30 μL twice | 0.627 | 18.93 |
| 9 | 30 μL twice | 0.609 | 18.48 |

### Functional annotation

The general functional analysis of the *de novo* assemblies and MAGs revealed a broad spectrum of COG (Clusters of Orthologous Genes) functional categories in our controlled and natural air samples. Briefly, gene predictions were made using Prodigal v2.6.3 ([ref](https://bmcbioinformatics.biomedcentral.com/articles/10.1186/1471-2105-11-119)), with COG functional categories analyzed using eggNOG v2.0.1 ([ref](https://academic.oup.com/nar/article/47/D1/D309/5173662)) and taxonomically classified using DIAMOND BLASTP. As eggNOG lacked taxonomic resolution, we also applied Prokka v1.14.6 ([ref](https://academic.oup.com/bioinformatics/article/30/14/2068/2390517)) followed by DIAMOND BLASTP to the bins, which delivered taxonomic and functional annotation. Following the findings in ‘Omics Insights in Environmental Bioremediation’, we filtered the annotated gene list to select genes involved in biodegradation and bioremediation. For comparing the functional inferences between the different sampling durations and locations, we calculated the relative abundance of the functional categories for the contigs (**Supplementary Figure 5A**) and MAGs (**Supplementary Figure 5B**) across samples of each experiment. We found a broad spectrum of genes encompassing diverse COG functional categories. The gene distribution was relatively similar between the controlled and the environmental setting, which was expected given the very basic metabolic and replication functionalities that are being described by COG.

The functional annotation of MAGs further allowed us to predict taxon-specific functions of the air microbiome. We, for example, obtained a *de novo* assembly of *Sphingomonas alba*, which has previously only been defined through a soil isolate (ref) and might therefore represent a novel strain with important functional variation. Our genome annotation identified genes (*flr, ribBA*) from flavin-based metabolic cycles, and a gene (*cher1*) which plays a role in biofilm formation and chemotaxis (ref). Certain bacterial taxa exhibit chemotactic responses towards aromatic hydrocarbons, which are prevalent pollutants, since they utilize these compounds as carbon sources; the *cher1* gene has been identified as a key gene in mediating this behavior (ref).

**Supplementary Figure 1.**
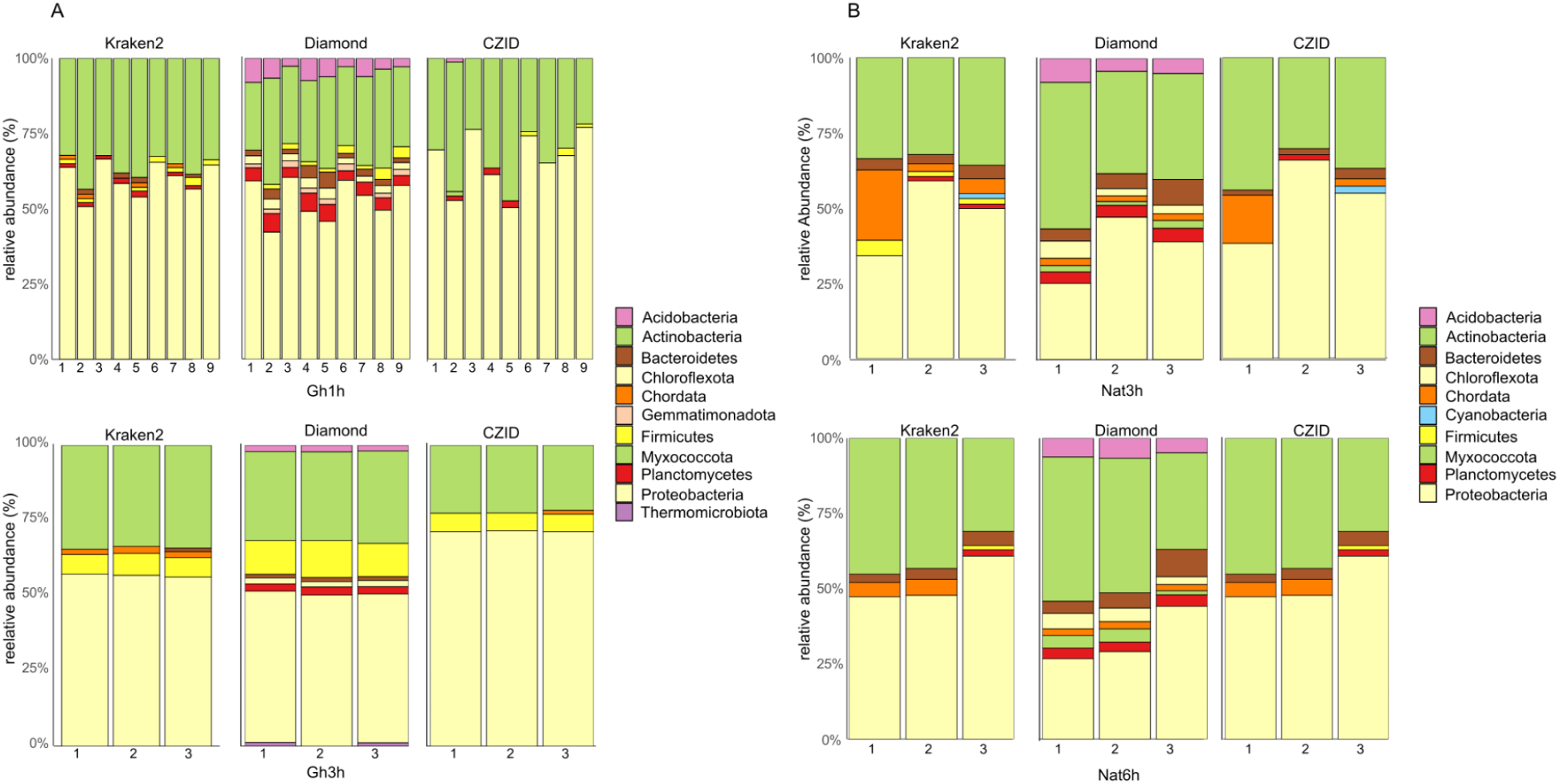
Taxonomic composition on the taxonomic phylum level using Kraken2, Diamond, and CZID annotations (Materials and Methods) of the **A**. controlled (Gh), and and **B**. natural (Nat) environment air samples.

**Supplementary Figure 2.**
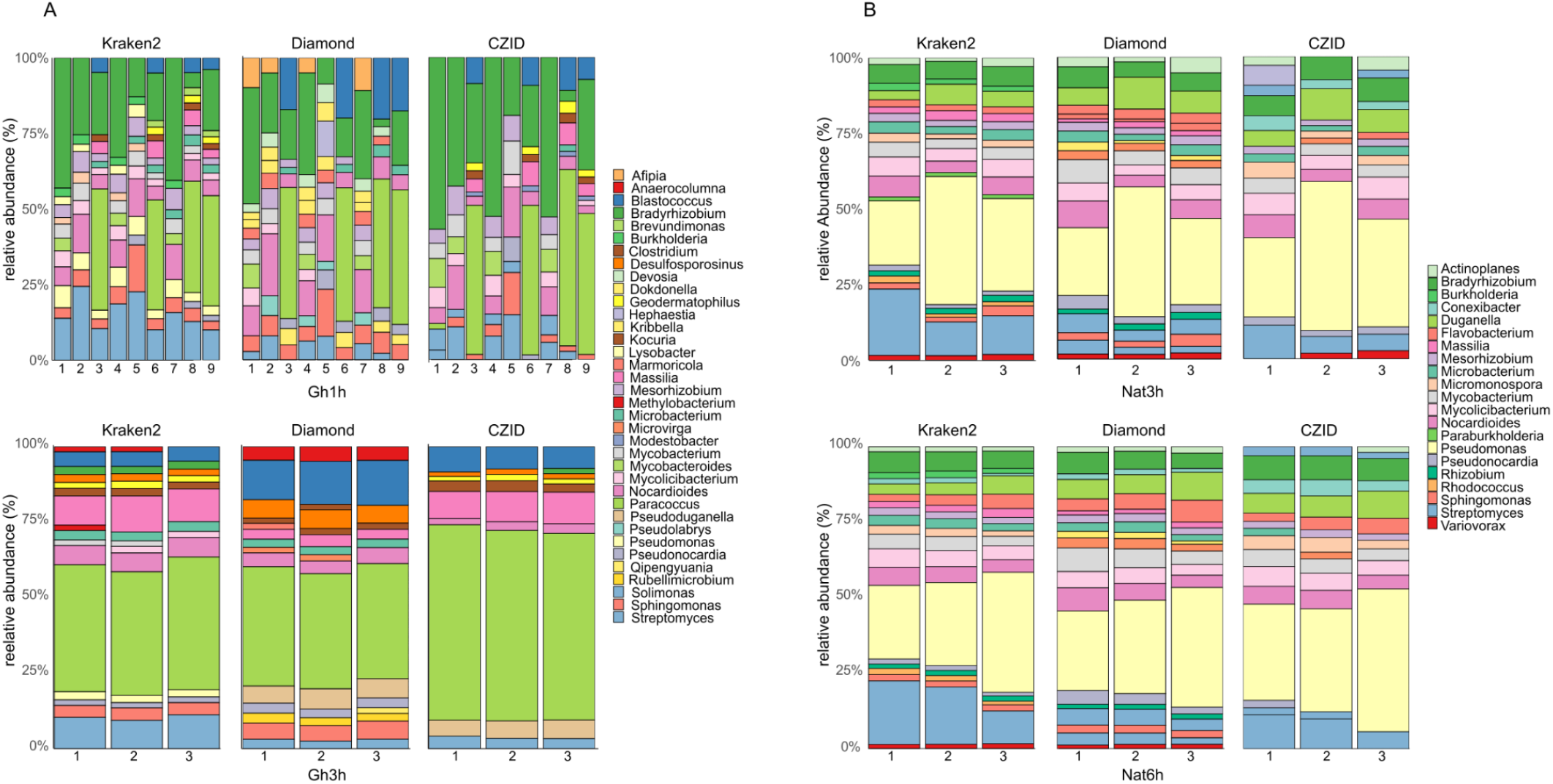
Taxonomic composition on the taxonomic genus level using Kraken2, Diamond, and CZID annotations (Materials and Methods) of the **A**. controlled (Gh), and and **B**. natural (Nat) environment air samples.

**Supplementary Figure 3.**
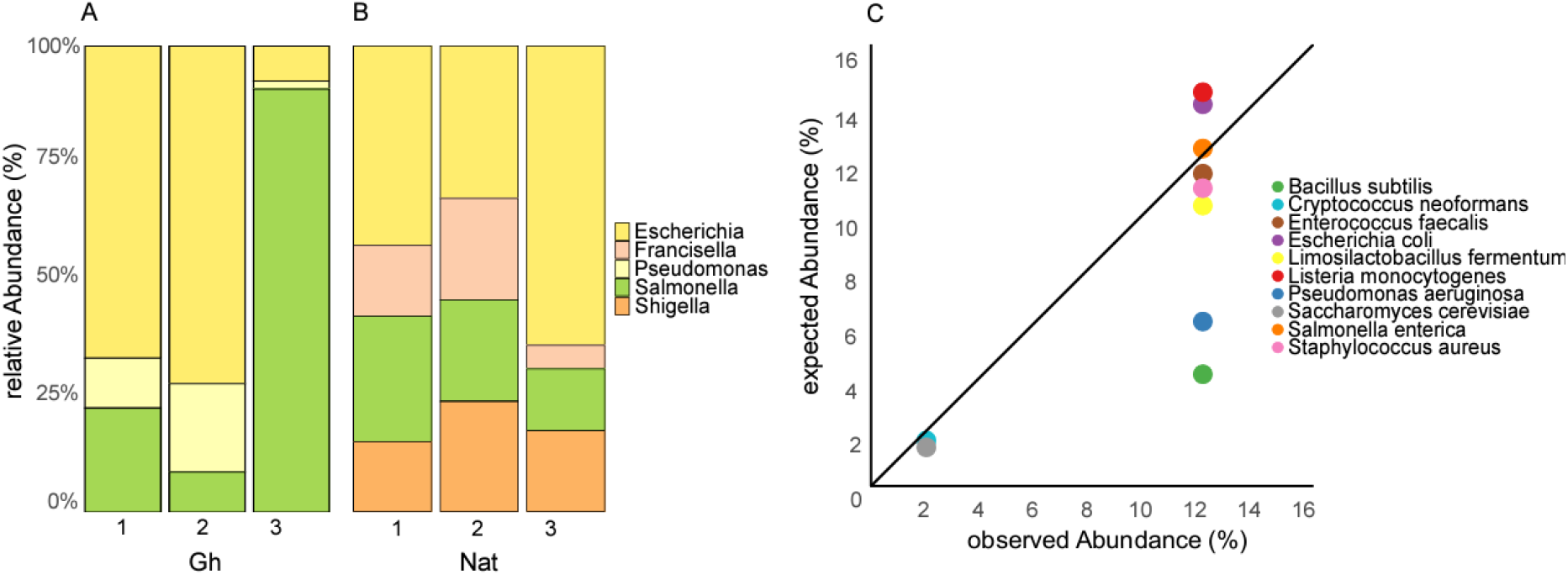
Relative abundance of microbial genera detected in the negative control samples from **A**. the controlled (Gh) and **B**. natural (Nat) environments (1: sampling control; 2: extraction control; 3: sequencing control). **C**. Relative expected versus observed relative abundance of microbial species in the positive control sample from a defined mock community (Materials and Methods).

**Supplementary Figure 4.**
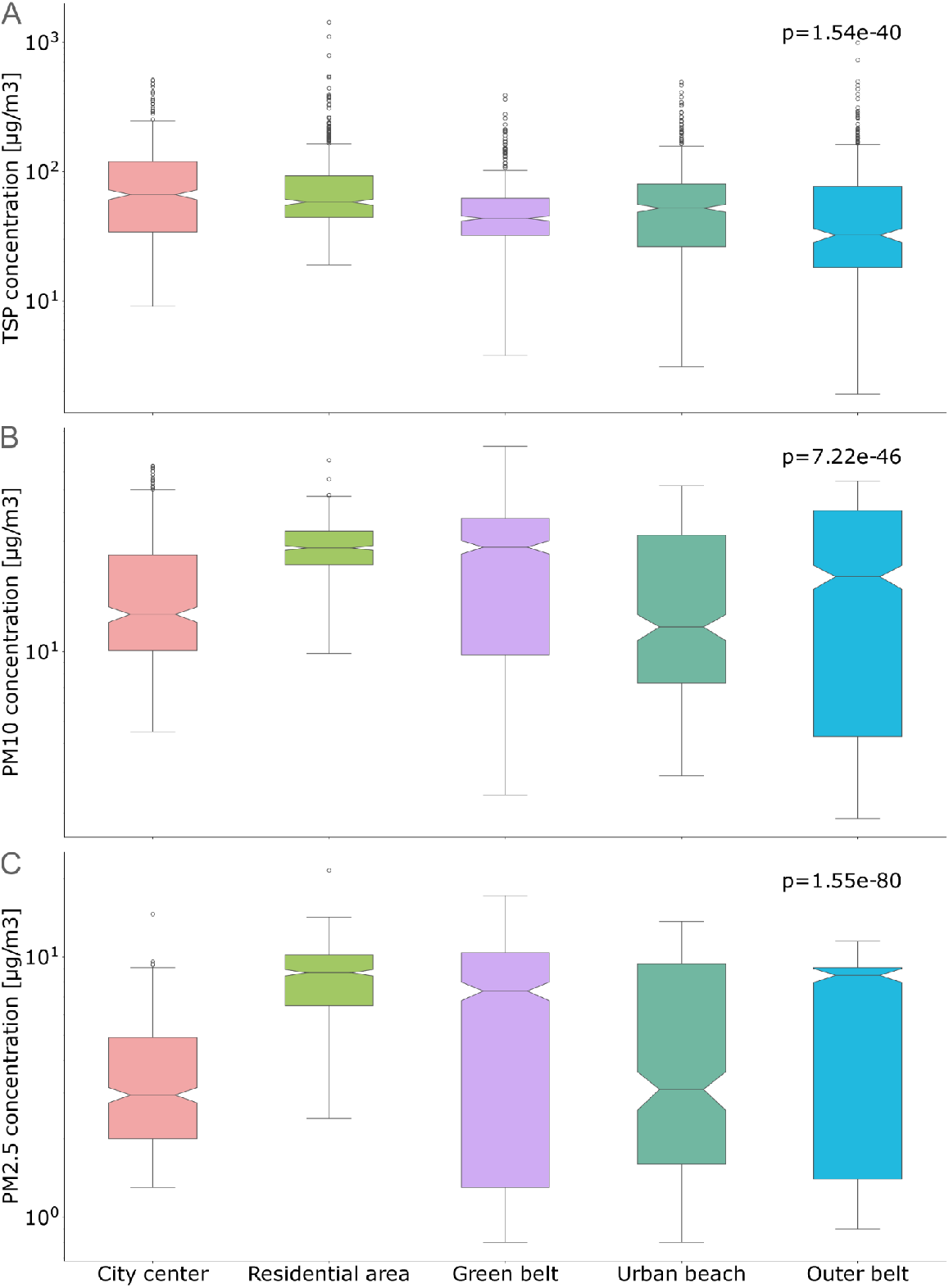
Particle mass fraction measurements across urban sampling locations for **A**. TSP, **B**. PM_10_, and **C**. PM_2.5_ [μg/m^3^]. The p-values describe the differences between all locations using the Kruskal-Wallis test (Materials and Methods).

**Supplementary Figure 5.**
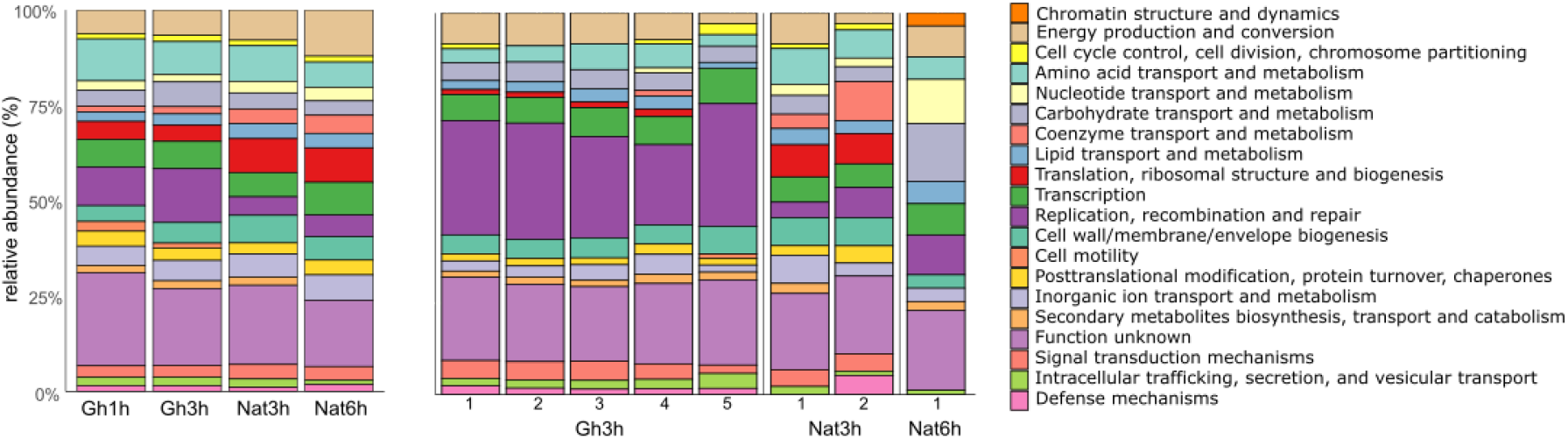
Analysis of COG functional categories in controlled (Gh) and natural (Nat) air samples. **A**. Annotation of assembled contigs. **B**. Annotation of MAGs. Each bar aggregates all functionalities detected across the respective samples from the same sampling condition within the same functional category.

